# Analysis of 26S Proteasome Activity Across Arabidopsis Tissues

**DOI:** 10.1101/2023.09.12.557456

**Authors:** Jagadeesan Ganapathy, Katherine A. Hand, Nitzan Shabek

**Affiliations:** Department of Plant Biology, College of Biological Sciences, University of California-Davis, Davis, CA 95616, USA

**Keywords:** 26S proteasome, plant proteolysis, cell-free system, fluorogenic probes, proteolytic activity, ubiquitin system, protein degradation, methods

## Abstract

Plants utilize the ubiquitin proteasome system (UPS) to orchestrate numerous essential cellular processes, including the rapid responses required to cope with abiotic and biotic stresses. The 26S proteasome serves as the central catalytic component of the UPS that allows for proteolytic degradation of ubiquitin-conjugated proteins in a highly specific manner. Despite the increasing number of studies employing cell-free degradation assays to dissect the pathways and target substrates of the UPS, the precise extraction methods of highly potent tissues remain unexplored. Here, we utilize a fluorogenic reporting assay using two extraction methods to survey proteasomal activity in different *Arabidopsis thaliana* tissues. This study provides new insights into the enrichment of activity and varied presence of proteasomes in specific plant tissues.

## Introduction

A major and essential protein-targeted degradation system known as the Ubiquitin Proteasome System (UPS) is involved in nearly every aspect of plant growth and development, including abiotic and biotic stress responses^1-6^. The targeted proteins are degraded through an ATP-dependent proteolysis process by the 26S proteasome, a massive (2.5MDa) multicatalytic protease complex^3,7-10^. Almost all targeted substrates of the proteasome are covalently labelled with polyubiquitin chains through a tightly regulated and highly specific cascade orchestrated by three enzymes: ubiquitin-activating enzyme E1, ubiquitin-conjugating enzyme E2, and a ubiquitin ligase E3^11,12^. Studies have shown that decreased 26S proteasome biogenesis in *Arabidopsis thaliana* leads to reduced cell division rates and heat shock hypersensitivity^13^. Dysfunction or variation in plant proteasomal activity and abundance also affected cell expansion and proliferation, as well as stress responses and crop performance^3,13^. Additional analyses of the proteasome subunit gene set demonstrated the highest expression of proteasomal genes in young organs and meristem tissues, implying that developmental stages need different, coordinated levels of activity^14-16^.

The 26S proteasome is composed of a 20S catalytic core particle (CP) and two 19S regulatory particles (RP)^3,7-10^. In particular, the 20S CP has a barrel shaped structure made up of four axially stacked heptameric rings (two outer α- and two inner β-rings) and catalyzes protein degradation^10,17,18^. More specifically, the inner β-rings contain seven different β-subunits; three of which (β1, β2 and β5) possess distinct proteolytic specificities: trypsin-like, chymotrypsin-like, and caspase-like activity, which allows for the cleavage of most peptide bonds^18,19^. The outer α-rings create a pore comprised of seven non-proteolytic α-subunits that allow for the entry and exit of target substrates and degradation products, respectively^20,21^. The 19S RP forms two sub-complexes known as the lid and base. These sub-complexes not only recognize specific polyubiquitinated proteins chains but also play roles in the deubiquitination, translocation, and unfolding of the targeted substrates^18,19^. Although the proteasome is highly conserved and essential, understanding of its regulation, activity, and abundance in different plant tissues is limited.

One traditional approach to measuring 26S proteasome activity utilizes short peptide substrates containing a C-terminal fluorophore, 7-Amino-4-methylcoumarin (AMC), such as succinyl-leucine-leucine-valine-tyrosine-AMC (Suc-LLVY-AMC)^22,23^. The method relies on proteolytic cleavage to liberate the AMC moiety from the peptide and measure fluorescence intensity^24^. This often requires successful isolation and purification of the proteasome, or the use of cell extracts enriched with proteasomes^24,25^. Increasing number of studies in planta have been employing cell-free degradation systems where plant cells extracts are utilized as native sources of the UPS machinery components^26,27^. This cell-free assay presents various challenges in determining the appropriate selection of specific plant tissues as well as successful extraction of a native and active UPS. Previous studies, for example, have revealed that starch and polyphenols frequently interfere with the isolation of plant proteins; further restricting the analysis of proteasome activity to specific tissues^25^. Despite the many advancements in the plant UPS field of study, the precise extraction methods of highly potent tissues that are enriched with active proteasomes remains to be fully explored. In this study, we surveyed 26S proteasomal activity in distinct *Arabidopsis thaliana* tissues, such as stems, roots, leaves, flowers, and seedlings, using two different lysis strategies. Our findings not only provide a benchmark strategy for a cell-free degradation system but also new insights into tissues enriched with proteasomal activity.

## Results and Discussion

To study the activity of the 26S proteasome in different parts of the plant, we first dissected the stems, roots, leaves, flowers, and seedlings of *Arabidopsis thaliana* that are known to highly express all proteasomal subunits (Fig. 1a-b). We then employed two distinct extraction methods, termed as Lysis 1 (L1, protein from frozen tissues that were extracted in the presence of liquid nitrogen using specific buffer, mortar and pestle) and Lysis 2 (L2, protein from frozen tissues that were extracted using native lysis buffer, plastic rod and a filter cartridge) (Fig. 1a, Fig. 2a, and Fig. S1a). The protein concentration was adjusted accordingly and normalized for each subjected sample from L1 and L2 lysates (Fig. 2a and Fig. S2c-d). Proteasomal activity was obtained in the presence and absence of proteasome inhibitor, MG132, and the fluorescence signal emitted by AMC cleavage was visualized and measured via a microplate reader (Fig. 1a and Fig. 2b-c). Interestingly, when utilizing L1 extraction method, significantly high levels of proteasomal activity were detected in the flowers, followed by the roots, leaves, seedlings, and stems (Fig. 2b). L2 also demonstrated the highest levels in the flowers, accompanied by the seedlings, leaves, stems, and roots (Fig. 2c). Markedly, no significant activity was measured in the roots (Fig. 2c), which may be attributed to the different extraction procedures. A comparison of several subunit genes belonging to the 19S RP and 20S CP exhibited the highest gene expression levels in the roots and flowers (Fig. 1b and Fig. S1b). This is comparable to our results, and we therefore propose that L1 is advantageous for deriving activity from the flowers, roots, and leaves, whereas L2 is more suitable for the seedlings and stems (Fig. 2d and Fig. S1). Furthermore, this data corroborates recent studies that employed cell-free proteasomal dependent degradation of recombinant proteins using native protein extracts from flowers^26-28^.

**Figure 1.**
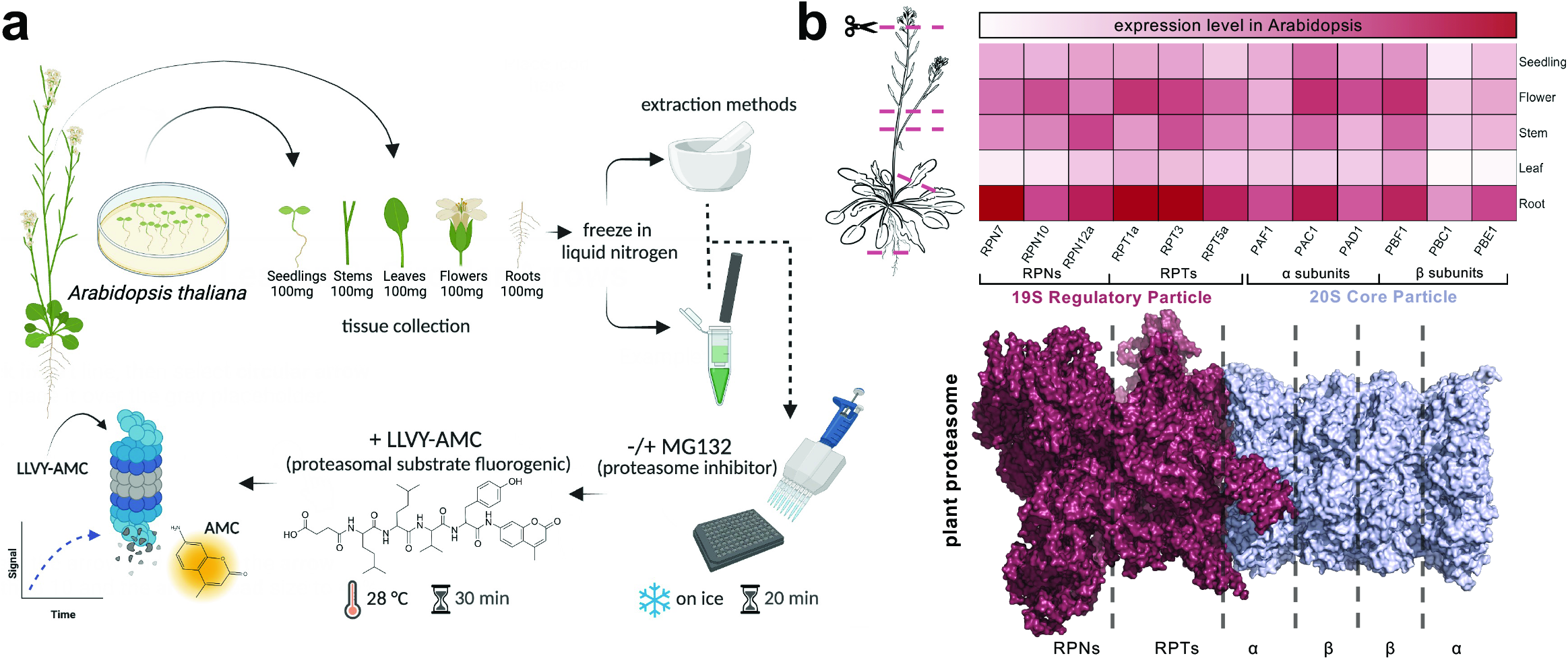
Schematic overview the 26S proteasome activity analysis across Arabidopsis tissues performed in this study. (a) Schematic representation of the proteasome activity assay using LLVY-AMC and the tissue sample preparation from *Arabidopsis thaliana* mature plant and seedlings using two different lysis strategies that were performed in this study. (b) Upper panel, heatmap comparison of several subunit genes from the 19S regulatory particle and 20S core particle in the seedlings, flower, stem, leaf, and root tissues of *Arabidopsis thaliana*. Data was obtained using the Klepikova Atlas from the Bio-Analytic Resource for Plant Biology and analyzed in R-Studio. Seedling expression combined and averaged the data from the seedling meristem, cotyledons, hypocotyl, and root. Highest levels of expression in the roots and flowers shown as darker red color denotes. Lower panel, architecture of plant proteasome from *Spinacia oleracea* (PDB: 7QVG and 7QVE)^34^. Graphical representation of the proteasome (shown as surface representation, 19S RP colored raspberry, 20S CP colored in light blue) was illustrated and analyzed via PyMOL-2.5.4. TAIR accession for the 19S RP presented are: *RPN7* (At4g24820), *RPN10* (At4g38630), *RPN12a* (At1g64520), *RPT1a* (At1g53750), *RPT3* (At5g58290), *RPT5a* (At3g05530). TAIR accession for the 20S CP presented are: *PAC1* (At3g22110), *PAD1* (At3g51260), *PAF1* (At5g42790), *PBC1* (At1g21720), *PBE1* (At1g13060), *PBF1* (At3g60820).

**Figure 2.**
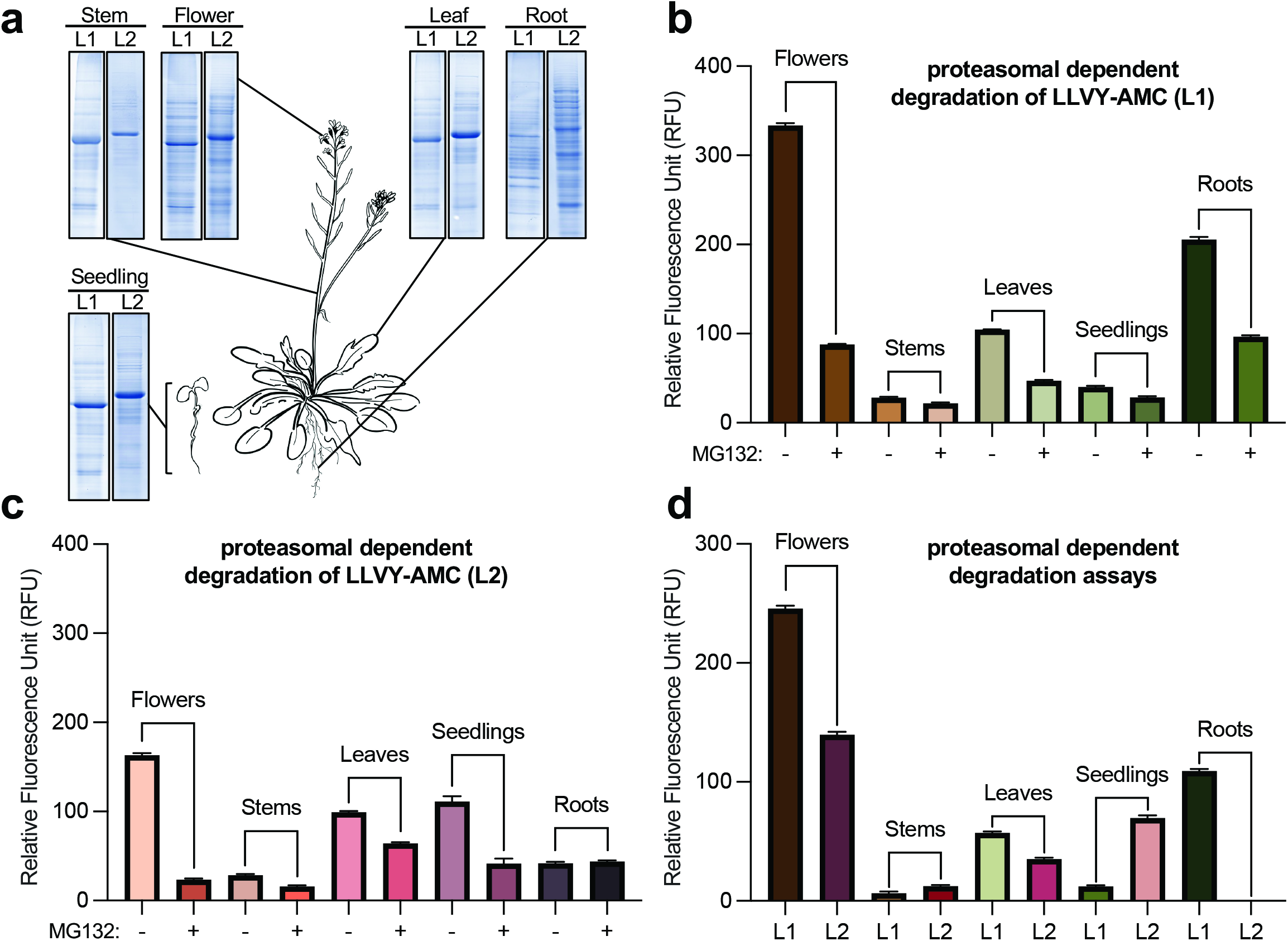
Proteasomal activity in different Arabidopsis tissues. (a) The dissected tissues that were used in the proteasomal activity assays were extracted and resolved via SDS-PAGE followed by Coomassie brilliant blue staining. *Arabidopsis thaliana* plant was drawn using Notability. (b-c) Proteasomal activity was measured in the presence and absence of proteasome inhibitor, MG132, for the different lysates 1 (L1, b) and 2 (L2, c) using the different tissues. (d) Comparative representation of the proteasomal activity-based assays shown in (c) and (d). The fluorescence signal emitted by AMC cleavage was visualized and measured via a microplate reader. The relative fluorescence unit (RFU) value measured via Gen5 software, averaged using Microsoft excel, and plotted using Graphpad Prism 10.0.1. Proteasome activity was plotted after subtracting the RFU value observed in the reactions without MG132. All experiments were repeated at least three times, and the plotted data shows the mean of three technical replicates. Error bars represent standard deviation (SD).

## Conclusion

Given its essential housekeeping role in catalyzing protein turnover events, the 26S proteasome provides cells with the proper machinery to alter their responses to the environment. Precise regulation, involving control of the quantity and presence of the proteasome, is crucial to ensure proper homeostasis in the cellular proteome as well. Recent findings revealed differences in proteasomal activity in mature and senescent leaves, and in developing siliques and seeds of *Arabidopsis thaliana*^*29*^. In this study, we observed clear distinctions in the abundance and activity of the 26S proteasome in the stems, roots, leaves, flowers, and seedlings of *Arabidopsis thaliana* between two lysis strategies. It is possible that higher proteasomal activity in tissues such as flowers and roots may be attributed to temporal developmental stages in plant growth. It has been previously demonstrated that the ubiquitin-proteasome system exhibits high activity in flowers, particularly when there is a pressing need for a rapid downregulation of specific proteins such as transcription factors^30-33^. Altogether, L1 and L2 are favorable extraction strategies for precisely investigating the activity and presence of the plant 26S proteasome in distinct tissues. Therefore, presenting suitable approaches to utilize highly potent tissues or cell types for targeted protein degradation by the proteasome both in vitro and in planta.

## Methods

### Materials and equipment

Magnesium chloride (MgCl_2_), glycerol, β-mercaptoethanol (β-ME), dithiothreitol (DTT), and MG-132 (Z-Leu-Leu-Leu-CHO) were purchased from Sigma-Aldrich (Saint Louis, MO, USA). Bioworld agar (phytoagar), Tris base, acetic acid, sodium hydroxide, magnesium chloride hexahydrate, Pierce™ protease inhibitor mini tablets (EDTA-free), ethanol, triton X-100, and the 96 well black/clear bottom plates were acquired from Thermo Fisher Scientific (Waltham, MA, USA). Adenosine 5’-triphosphate disodium salt hydrate (ATP) was purchased from Research Products International (Mount Prospect, IL, USA). Suc-LLVY-AMC (chymotrypsin-like activity substrate) was obtained from Cayman Chemicals (Ann Arbor, MI, USA). Minute™ total protein extraction kit and Minute™ native lysis buffer (used for L2 method) were purchased from Invent Biotechnologies (Plymouth, MN, USA). Bio-Safe™ Coomassie stain and Precision Plus Protein™ dual color standard were obtained from Bio-Rad (Hercules, CA, USA). Gamborg’s B-5 Medium was acquired from Life Technologies (Carlsbad, CA, USA). The substrate solution is prepared by dissolving Suc-LLVY-AMC in DMSO to a final concentration of 10mM. The proteasome inhibitor MG132 was dissolved in DMSO to a final concentration of 10mM. Lysis buffer 1 (L1) contains 50 mM Tris-HCl pH 7.4, 10% glycerol, 0.01% triton X-100, 5mM β-ME and protease inhibitor cocktails. The assay master mix buffer has 50 mM Tris-HCl pH 7.4, 10 mM MgCl2, 1 mM ATP, and 1 mM DTT.

The equipment used in this study included a pH meter (Hanna Instruments), Barnstead Labquake Shaker Rotisserie (The Lab World Group), protein electrophoresis equipment (Bio-Rad), ChemiDoc Touch Imaging system (Bio-Rad), microcentrifuge (Beckman Coulter), VWR 1500E Incubator (Marshall Scientific), Laminar Flow Hood (Labconco), and plate reader spectrophotometer (BioTek Instruments, model Synergy H1).

### Plant growth conditions

For flower, stem, and leaf analysis, *Arabidopsis thaliana* (Col-0 background) seeds were grown on Sunshine Mix #1 Fafard 1P (Sungro Horticulture, Agawam, MA, USA) soil in a growth chamber with a 16-hour light/8-hour dark photoperiod. For seedling and root tissue analysis, seeds were sterilized with 70% EtOH, plated on agar medium supplied with B5 media, and grown in a growth chamber with a 16-hour light/8-hour dark photoperiod.

### Plant tissue preparation

100 mg of leaves, stems, flowers, and roots were harvested after 1 month and 5 days, and seedling tissues were collected after 12 days, wrapped in aluminum foil, and frozen in liquid nitrogen. For extraction method I (L1), Total proteins were extracted by mechanically lysing 100 mg of each tissue in 100 μL of L1 buffer using a mortar and pestle, and incubating for 5 minutes at 4°C. Following incubation, tissue extracts were centrifuged at 15,000 RPM for 5 minutes at 4°C. The supernatant was collected, and the extract concentrations were measured using the Bradford method (Bio-Rad Bradford reagent). All sample extracts were adjusted to 1 mg/mL. For extraction method II (L2), total proteins were extracted by combining 100 mg of each tissue with 100 μL of Minute™ native buffer, lysing with plastic rods, and following the manufacturer’s instructions. Lysates (L1 and L2) were incubated for 5 minutes at 4°C and centrifuged at 15,000 RPM for 5 minutes at 4°C. The supernatants were recovered, and the concentrations were measured using the Bradford method. All extracts were adjusted to 1 mg/mL in the final reactions (supplemented with protease inhibitors cocktail). We found that long-term storage of extract samples is not recommended as it may decrease proteasome activity, therefore all experiments were done up to a week following the extraction protocols.

### Proteasome activity assay

Proteasome activity assays were performed in reaction mixture (1 mg/mL tissue extract, 20 μM LLVY-AMC (substrate), -/+ 100 μM MG132 (proteasome inhibitor), 18.5 μL of assay master mix buffer, and 1% SDS) in a 50 μL volume on a 96-well plate. All assays were performed in triplicate reactions. 1 mg/mL of tissue extract was mixed with 18.5 μL of assay master mix buffer in the presence or absence of 100 μM MG132 in a 96-well plate, incubated on ice for 20 minutes, and then supplemented with 20 μM LLVY-AMC. The plate was incubated for 30 minutes at 28°C and the reactions were terminated by the addition of 1% SDS. For control, similar assay master mix buffer and extraction buffer were added to one well, and another well contained only tissue extract and assay master mix buffer, all in equivalent volumes with or without LLVY-AMC. 26S proteasome activity was measured by a Synergy H1 Microplate Reader with excitation of 350 nm and emission of 440 nm using 1-minute intervals over 5-minutes. The relative fluorescence unit (RFU) value measured via Gen5 software, averaged using Microsoft excel, and plotted using Graphpad Prism 10.0.1. Standard deviations were calculated and presented as error bars together with the mean values. Proteasome activity was plotted after subtracting the RFU value observed in the reactions without MG132.

## Acknowledgement

N.S. is supported by the National Science Foundation (NSF-CAREER Award #2047396, NSF-EAGER Award #2028283, and Award #2139805), and by the U.S. Department of Energy, Office of Science, Biological and Environmental Research, Genomic Science Program grant no. DE-SC0023158.

## Author Contribution

J.G. and N.S. conceived and designed the experiments. J.G. conducted the protein extractions and proteasome activity assays. J.G., K.A.H., and N.S. analyzed the proteasome activity assays. N.S. and K.A.H. wrote the manuscript with input from J.G.

## Data Availability

All relevant data are available from the corresponding author upon reasonable request.

