## Supplemental Figures for "Analysis of 26S Proteasome Activity Across Arabidopsis Tissues"

#### Supplementary Figures

**Figure S1**: Extended schematic representation of the fluorogenic activity assay using two different extraction methods analyzed in this study and 26S proteasome subunit gene expression in different tissues of *Arabidopsis thaliana.*

**Figure. S2:** Proteasome dependent activity in each tissue of *Arabidopsis thaliana* for L1 and L2 strategies.

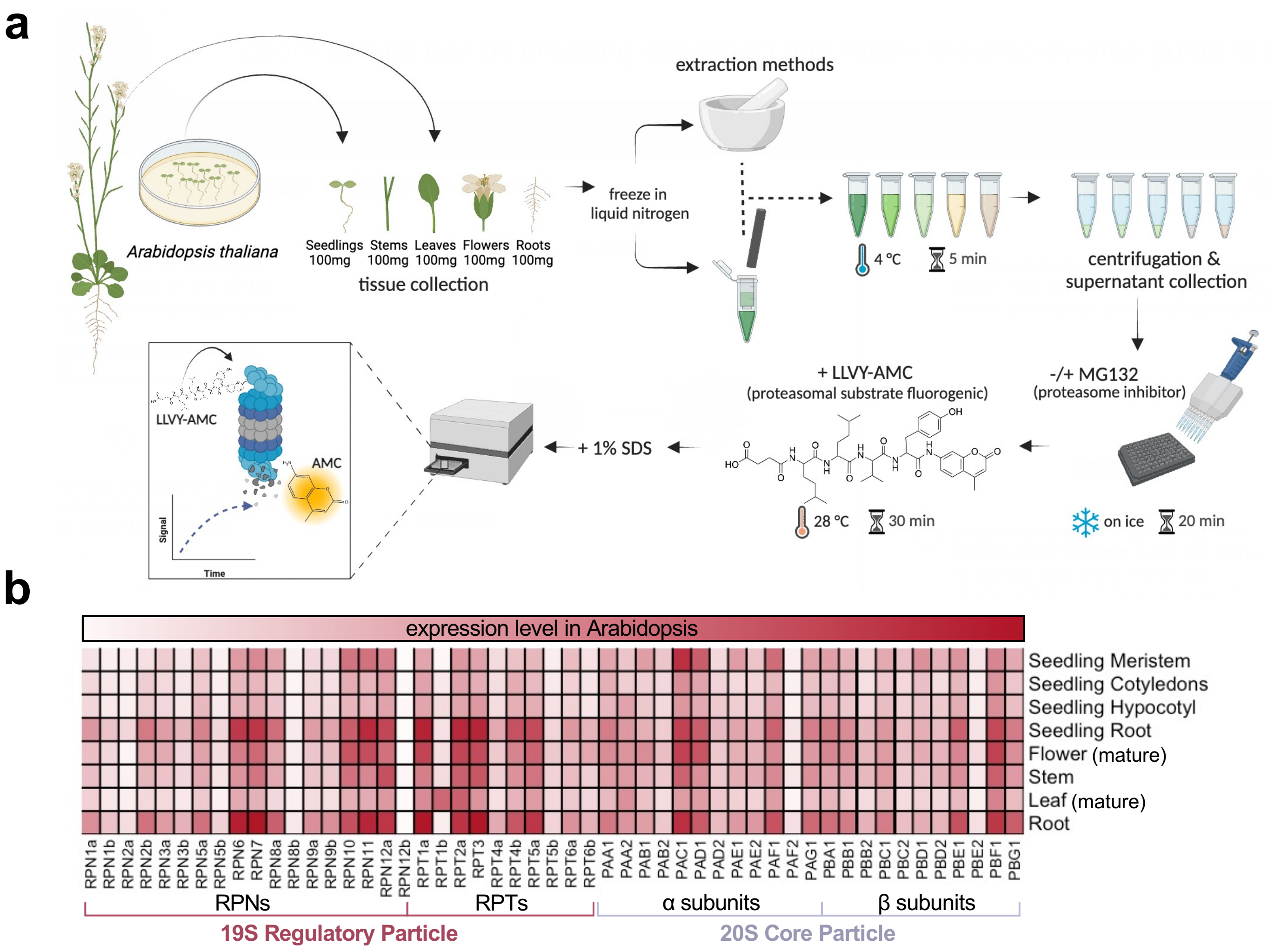


**Supplementary Figure 1.** (a) Extended schematic representation of the fluorogenic activity assay using two different extraction methods analyzed in this study. (b) Extended heatmap demonstrating 26S proteasome subunit gene expression in different tissues of *Arabidopsis thaliana*. Data was obtained using the Klepikova Atlas from the Bio-Analytic Resource for Plant Biology and analyzed in R-Studio. Darker red color denotes higher gene expression. TAIR accession for the 19S RP presented are: *RPN1a* (At2g20580), *RPN1b* (At4g28470), *RPN2a* (At1g08410), *RPN2b* (At2g32730), *RPN3a* (At1g20200), *RPN3b* (At1g75990), *RPN5a* (At5g09900), *RPN5b* (At5g64760), *RPN6* (At1g29150), *RPN7* (At4g24820), *RPN8a* (At5g05780), *RPN8b* (At3g11270), *RPN9a* (At5g45620), *RPN9b* (At4g19006), *RPN10* (At4g38630), *RPN11* (At5g23540), *RPN12a* (At1g64520), *RPN12b* (At5g42040), *RPT1a* (At1g53750), *RPT1b* (At1g53780), *RPT2a* (At4g29040), *RPT3* (At5g58290), *RPT4a* (At5g43010), *RPT4b* (At1g45000), *RPT5a* (At3g05530), *RPT5b* (At1g09100), *RPT6a* (At5g19990), *RPT6b* (At5g20000). TAIR accession for the 20S CP presented are: *PAA1* (At5g35590), *PAA2* (At2g05840), *PAB1* (At1g16470), *PAB2* (At1g79210), *PAC1* (At3g22110), *PAD1* (At3g51260), *PAD2* (At5g66140), *PAE1* (At1g53850), *PAE2* (At3g14290), *PAF1* (At5g42790), *PAF2* (At1g47250), *PAG1* (At2g27020), *PBA1* (At4g31300), *PBB1* (At3g27430), *PBB2* (At5g40580), *PBC1* (At1g21720), *PBC2* (At1g77440), *PBD1* (At3g22630), *PBD2* (At4g14800), *PBE1* (At1g13060), *PBE2* (At3g26340), *PBF1* (At3g60820), *PBG1* (At1g56450).

### Supplementary Figure 1

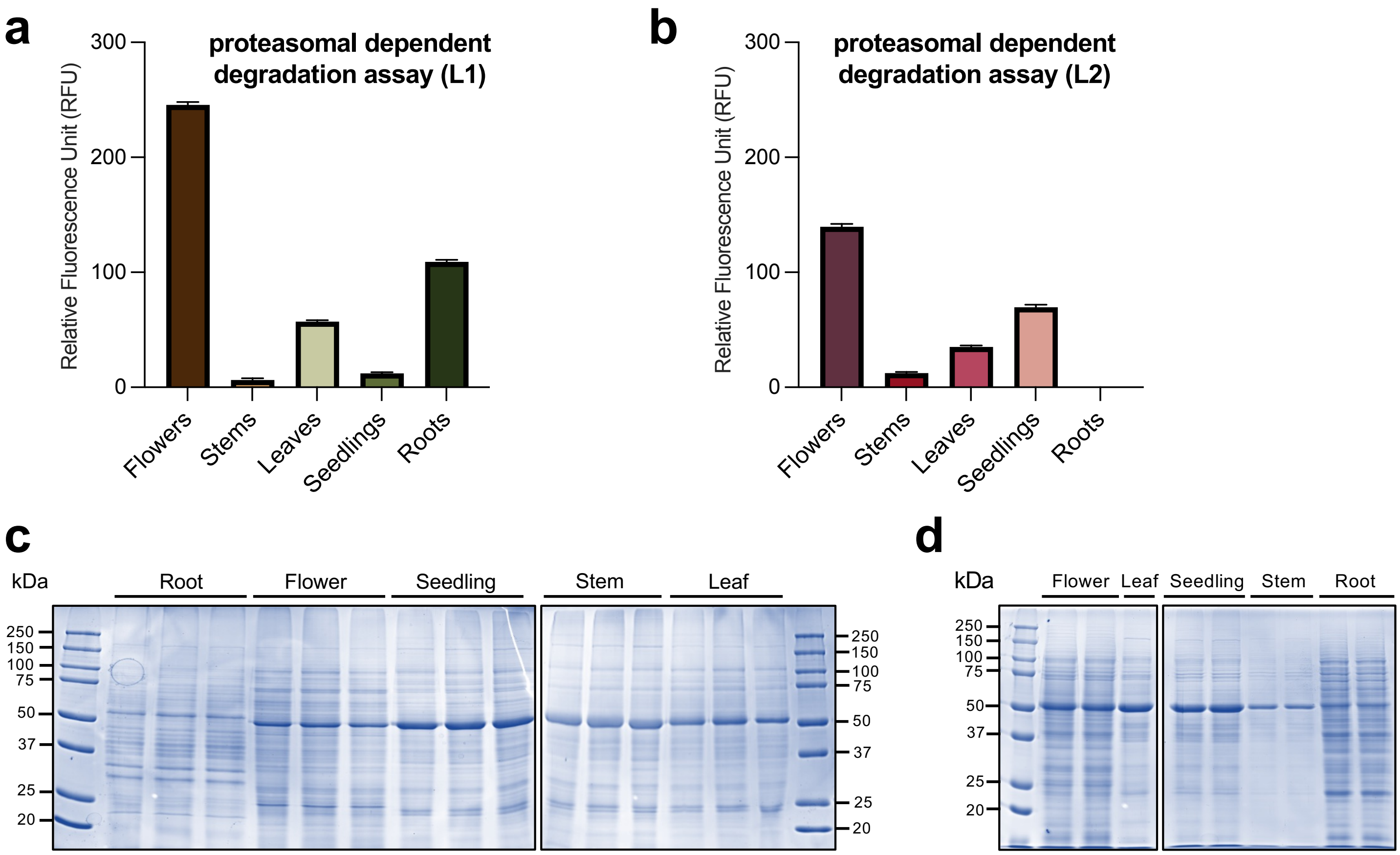


**Supplementary Figure 2.** Proteasome dependent activity in each tissue of Arabidopsis thaliana for L1 and L2. (a-b) Data in the absence of MG132 was subtracted from the data in the presence of MG132 for each tissue extract. The relative fluorescence unit (RFU) value measured via Gen5 software, averaged using Microsoft excel, and plotted using Graphpad Prism 10.0.1. Proteasome activity was plotted after subtracting the RFU value observed in the reactions without MG132. All experiments were repeated at least three times and the plotted shows the mean of three technical replicates. Error bars represent standard deviation (SD). (c-d) Represented reaction mixtures containing extracts from the different tissues used in this study were resolved via SDS-PAGE and visualized by Coomassie staining.

### Supplementary Figure 2
